# Genomics in healthcare: GA4GH looks to 2022

**DOI:** 10.1101/203554

**Authors:** Ewan Birney, Jessica Vamathevan, Peter Goodhand

**Author notes:** Correspondence to: Ewan Birney, EMBL-EBI, Wellcome Genome Campus, Hinxton, Cambridge CB10 1SD, United Kingdom.

## Abstract

The Global Alliance for Genomics and Health (GA4GH), the standards-setting body in genomics for healthcare, aims to accelerate biomedical advancement globally. We describe the differences between healthcare- and research-driven genomics, discuss the implications of global, population-scale collections of human data for research, and outline mission-critical considerations in ethics, regulation, technology, data protection, and society. We present a crude model for estimating the rate of healthcare-funded genomes worldwide that accounts for the preparedness of each country for genomics, and infers a progression of cancer-related sequencing over time. We estimate that over 60 million patients will have their genome sequenced in a healthcare context by 2025. This represents a large technical challenge for healthcare systems, and a huge opportunity for research. We identify eight major practical, principled arguments to support the position that virtual cohorts of 100 million people or more would have tangible research benefits.

## Introduction

Human genomics^1^ is undergoing a shift from a predominantly research-driven activity to one that is driven—and funded—by health care. Human datasets collected by healthcare providers and made available to researchers now offer unprecedented opportunities for rapid advancement of biological research. Humans have always been a focus for research, both in the clinic and in probing fundamental life processes, but the scale and scope of these efforts have been relatively modest. Healthcare funding for genomic sequencing of humans on a population scale is providing remarkable, transformative opportunities for both clinical practise and basic research. This is the premise of the Global Alliance for Genomics and Health (GA4GH), which seeks to facilitate the coalescence of clinical practise and biomedical research for mutual benefit to clinicians, researchers, and human health.

It is an exciting time for genomics researchers because results from our work can now have a direct impact on healthcare—a major goal for many researchers. This change is driven by the growing effectiveness of genomics in practising medicine and by the increasing affordability of genomics technology. Rolling out genomics in healthcare will open up opportunities to study the genetic and molecular components of health and disease on an unprecedented scale. But the challenge of transferring expertise effectively from the research domain to healthcare is massive, rivalled only by that of establishing data-access mechanisms that are both appropriate to research applications and respectful of the rights of the individuals to whom the data pertain. We believe these challenges can be met, but only if the genomics community is committed to broad-based advocacy and coordinated efforts worldwide.

This is not a new position. The community has anticipated this shift since the Human Genome Project began in the late 1980s, and the trends we summarise here have been the subject of keen observation for many years. The current transition is a significant moment for genomics and healthcare; but there have been step changes along the way, and many more are still to come. Here, we simply pause to reflect on the landscape of genomics—to consider the future paths before us and the role GA4GH can play in accelerating biomedical advancement.

### The Global Alliance for Genomics and Health

GA4GH is a worldwide alliance of genomics and clinical researchers, data scientists, healthcare practitioners, and many others working together to establish frameworks and standards for responsible, international genomic and health-related data sharing. Our ultimate goal is to leverage the detailed knowledge of the genomics research community to advance healthcare and, conversely, to enable the enrichment of biomedical research through the responsible sharing and use of clinical genomic data in research. It is clear that without such a consortium, the uptake of genomics into clinical practise will be slower, more expensive and riskier, and will differ country by country with little harmonisation. This would reduce the benefit to patients worldwide substantially, and increase costs to healthcare systems. In such a landscape, we would not be able to tap into the opportunities presented by millions of human genomic sequence and phenotypic data for basic research.

Through this lens, we describe the differences between healthcare- and research-driven genomics, discuss the implications of global, population-scale collections of human data for research, and outline mission-critical considerations in ethics, regulation, technology, data protection, and society.

## Genomics in healthcare

The process of sequencing a genome is essentially the same in any setting, but the scale of production and regulation of the resulting data are quite different when used for health versus research. For many research projects, the cost of sequencing is a large—if not the largest—line item in the budget. ‘Research genomes’ are, by convention, open to other researchers following publication (if not before), often with managed access (via the database of Genotypes and Phenotypes, dbGaP or the European Genome-phenome Archive, EGA) to ensure compliance with the consent structure the patients have agreed to for research use of their data. It is commonplace for researchers worldwide to draw upon such datasets from a variety of studies, which increases the amount of knowledge derived from each genome.

Healthcare has an entirely different financial, legal, and social landscape (see Table 1). It is the largest component of the economies of the G20 countries, and one of the most regulated industries. Its structure and regulation are very diverse from country to country, covering the full spectrum from state-run to private provision. In each system, the cost of an assay in healthcare—genomics included—is always considered in light of its benefits to the health of an individual. In theory, if a genome assay becomes cost–effective for a specific application within a healthcare system, the only limit to its deployment is the number of patients who will potentially benefit via that application. In practise, there are many logistical and regulatory hurdles to overcome before an assay can be incorporated into a healthcare system’s regular offerings.

**Table 1.** Differences between genomics for research and genomics for healthcare

|  | Research Genomics | Healthcare Genomics |
| --- | --- | --- |
| **Rationale for genome sequencing** | Ability to answer research study questions | Ability to impact healthcare diagnosis and treatment decisions of an individual |
| **Number of genomes sequences needed** | Dependent on power analysis to observe desired effects (in practise, genome sequencing cost often limiting) | Dependent on population size of patients with specific criteria (in theory, independent of patient population size once shown to be cost effective) |
| **Desired timelines of sequencing and analysis** | Up to 5 years, can be far longer | Weeks to months depending on disease |
| **Data sharing to researchers** | Routine after publication, occasionally before | Varies by system, but usually not outside a joint research component |
| **Legal framework for sharing data** | Broadly based around ethics and data access committees set up for research | Varies nation by nation, usually with explicit legislative mechanisms |
| **Language for agreements and discussion** | English as international language of science | National language(s) |
| **Estimated data size by 2025** | ~Petabytes to low Exabytes | Exabytes to low Zetabytes worldwide |
| **Data sharing mechanisms** | Aggregation of anonymised data, and local download & processing | Federated access to secure data to virtual/containerised analysis, pooling of summary/intermediate statistics |

The current case for implementing genomics in healthcare can be presented in at least four broad categories: infectious disease, rare disease, cancer, and common or chronic disease. These categories map to clear clinical use cases, though there is—as always in human biology—huge overlap between them. For example, there is a continuum between rare and common disease, and rare disease and severe cancer susceptibility can both be diagnosed and managed following similar clinical genetic pathways. Infectious disease contributes to both cancer and chronic disease, often as a key trigger.

Despite the inevitable crossover, these four categories map well to the funding and management approaches of many different healthcare systems. In the following sections, we outline the case for genomics in each category.

### Infectious disease

Genomics can be used to identify the infectious agent of disease with more confidence and precision than ever before, and at increasing speed (1,2). The main challenges to deployment in healthcare are managing cost and logistics, achieving precise phenotypic prediction (e.g. antibiotic resistance), and aligning historical records of non-genomic-based assays with contemporary genomics tests.

Genomic sequencing of pathogens for diagnosing infectious disease is quickly ramping up within healthcare systems around the world. Recent announcements (3) for large-scale tuberculosis sequencing in several countries, rapid identification of Ebola and Zika virus strains (4), and tracing hospital outbreaks using genomics (5,6) all demonstrate a vibrant, functional interface between research and clinical practise.

### Rare disease

Rare diseases (i.e. those with a person frequency of 1 in 2,000 or lower (7)) often have a clear genetic component, often of high penetrance with only a few genes involved in each disease. Clinical geneticists have used single-gene tests since the early 1990s to support diagnosis and some treatment decisions for many of these diseases.

Genomics provides a confident diagnosis for many rare diseases, which enables families and healthcare systems to manage the disease appropriately and brings patient retesting (a.k.a. their “diagnostic odyssey”) to a close. Genomic diagnosis of a child can inform parents about the odds of the disease affecting a future child (i.e. distinguishing de novo mutations from recessive alleles), empowering them to make informed family-planning decisions. In exceptional cases, a successful diagnosis can lead to profound changes in a patient’s medical treatment, prognosis, and quality of life.

The cost of assaying broader genomic regions, including whole-genome sequencing, has fallen considerably, which has had a substantial impact on rare-disease diagnosis and research (8,9). Arguably, the rare-disease space has seen the most successful deployment of genomics in healthcare, with multiple systems reporting diagnostic rates of between 20% and 30%, and specific health economics papers providing multiple lines of evidence of cost-effectiveness (10).

### Cancer

One of the consistent hallmarks of cancer is altered somatic genomes, often with specific mutations that ‘drive’ the cancer. Characterising a cancer by sequencing a tumour’s genome alongside the patient’s unaffected genome has returned profound insights for cancer research.

Applying cancer genomics in the clinic is more complicated. The time window for care decisions is measured in weeks rather than months, and gaining genomic information on such a short timescale is logistically challenging, to say the least. In addition, while straightforward diagnosis has immediate benefits for rare-disease patients, cancer genomic information is only considered useful if it changes treatment options. For this to occur, carefully planned clinical trials are necessary to chart change of treatment impact on the basis of genomics.

The heterogeneity of cancer as a disease—of each individual cancer and of any subsequent metastasis—adds many layers of complexity to genomic analysis. However, oncologists are increasingly confident that genomic information will be useful in cancer care decisionmaking. Indeed, it is being deployed effectively in a growing number of cases, and we are confident that the roll-out of genomics into practicing cancer care will increase steadily in the coming years.

### Common or chronic disease

‘Common disease’ is a catchall phrase describing a vast spectrum of diseases that have distinct ‘environmental’ and ‘genetic’ components. This area of genomics is firmly rooted in research, and is the healthcare category where genomic information is most removed from practise, even though there is considerable effort on the research use of genomics.

The focus of genomic research in common disease is large-scale cohorts, for example biobanks in the UK (UK BioBank, Generation Scotland), China (Kadoorie), and the US (the nascent All of Us Research Program); megabanks such as Tohoku in Japan; and the whole-population cohorts in Iceland, Estonia, Finland (the Sisu project), and Denmark. Genotyping has become a regular feature of large-scale cohorts, and whole-genome sequencing is bound to become routine here as well.

Currently, the value of genomics for common diseases such as cardiovascular disease and diabetes is mainly in research. However, some clinical practise work is being explored, for example adapting screening protocols (11).

## How many genomes?

The four categories of healthcare we describe above differ in their potential penetration into the population and in their direct involvement of individual people.

**Infectious-disease** genomics currently focuses on sequencing infectious agents, and as such contributes huge amounts of genomic information on bacteria and viruses, but little on humans.

**Rare diseases** affect up to 1 in 7 (14%) people in G20 countries. Restricting this figure to the numbers projected in countries with active genomic rare disease diagnosis processes, the percentage of people affected is closer to 1% to 2% of live births (depending on the healthcare system). Both parents of the patient are sequenced whenever possible, so in countries where rare-disease genomics is deployed in healthcare, between 2% and 5% of the population could be participating actively in this category of genomic testing. Given the robust benefit to healthcare, logistical simplicity, and cost-effectiveness compared to current approaches, at the minimum prevalence of 1% of the population, 20 million people could well be sequenced for rare-disease diagnosis by 2025 (see Table 2).

**Table 2.**
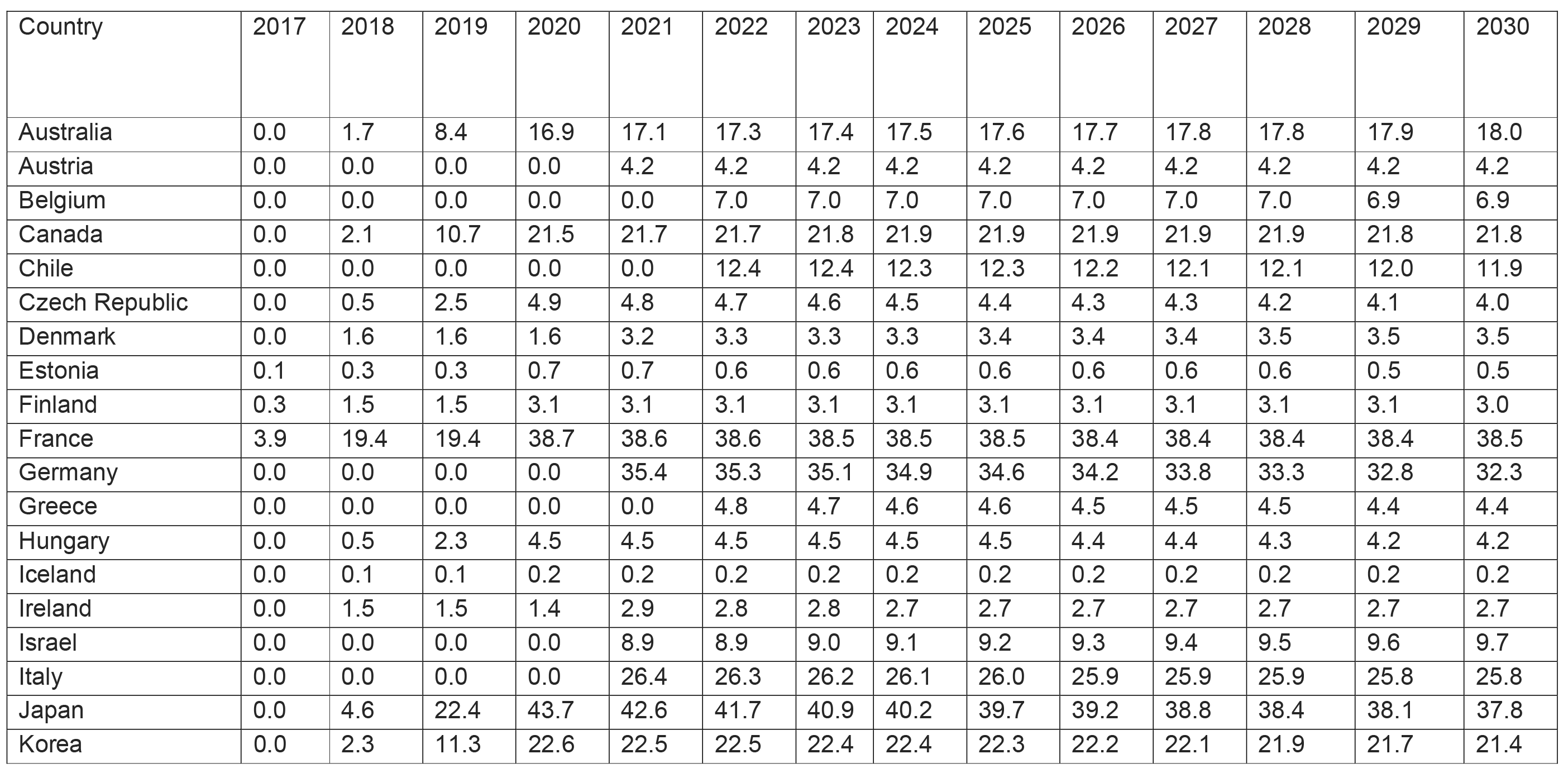
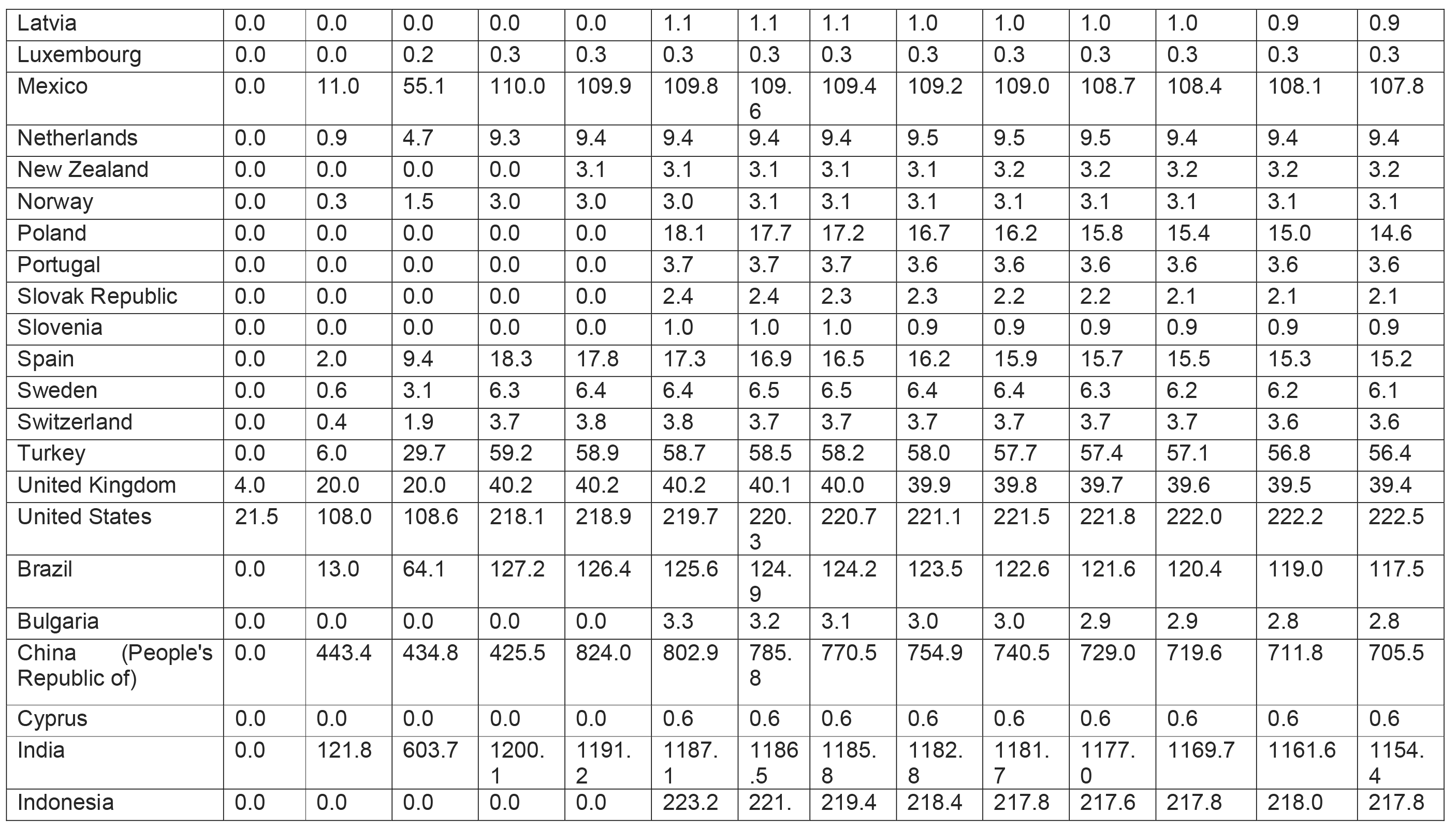
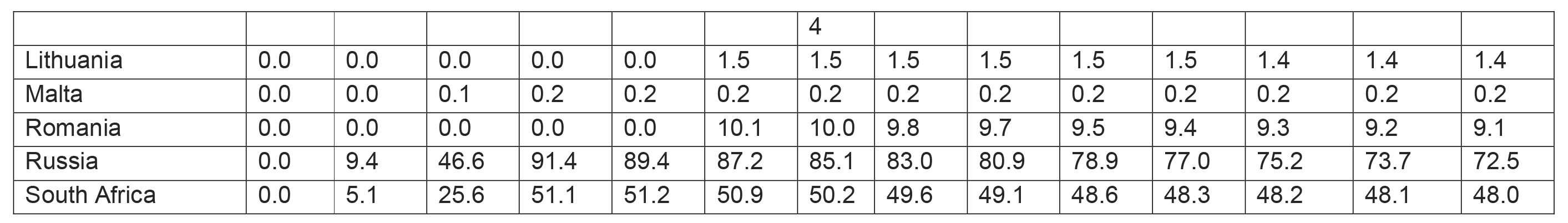
Estimated number of Rare disease genomes (in thousands) sequenced in each OECD country over time

| Country | 2017 | 2018 | 2019 | 2020 | 2021 | 2022 | 2023 | 2024 | 2025 | 2026 | 2027 | 2028 | 2029 | 2030 |
| --- | --- | --- | --- | --- | --- | --- | --- | --- | --- | --- | --- | --- | --- | --- |
| Australia | 0.0 | 1.7 | 8.4 | 16.9 | 17.1 | 17.3 | 17.4 | 17.5 | 17.6 | 17.7 | 17.8 | 17.8 | 17.9 | 18.0 |
| Austria | 0.0 | 0.0 | 0.0 | 0.0 | 4.2 | 4.2 | 4.2 | 4.2 | 4.2 | 4.2 | 4.2 | 4.2 | 4.2 | 4.2 |
| Belgium | 0.0 | 0.0 | 0.0 | 0.0 | 0.0 | 7.0 | 7.0 | 7.0 | 7.0 | 7.0 | 7.0 | 7.0 | 6.9 | 6.9 |
| Canada | 0.0 | 2.1 | 10.7 | 21.5 | 21.7 | 21.7 | 21.8 | 21.9 | 21.9 | 21.9 | 21.9 | 21.9 | 21.8 | 21.8 |
| Chile | 0.0 | 0.0 | 0.0 | 0.0 | 0.0 | 12.4 | 12.4 | 12.3 | 12.3 | 12.2 | 12.1 | 12.1 | 12.0 | 11.9 |
| Czech Republic | 0.0 | 0.5 | 2.5 | 4.9 | 4.8 | 4.7 | 4.6 | 4.5 | 4.4 | 4.3 | 4.3 | 4.2 | 4.1 | 4.0 |
| Denmark | 0.0 | 1.6 | 1.6 | 1.6 | 3.2 | 3.3 | 3.3 | 3.3 | 3.4 | 3.4 | 3.4 | 3.5 | 3.5 | 3.5 |
| Estonia | 0.1 | 0.3 | 0.3 | 0.7 | 0.7 | 0.6 | 0.6 | 0.6 | 0.6 | 0.6 | 0.6 | 0.6 | 0.5 | 0.5 |
| Finland | 0.3 | 1.5 | 1.5 | 3.1 | 3.1 | 3.1 | 3.1 | 3.1 | 3.1 | 3.1 | 3.1 | 3.1 | 3.1 | 3.0 |
| France | 3.9 | 19.4 | 19.4 | 38.7 | 38.6 | 38.6 | 38.5 | 38.5 | 38.5 | 38.4 | 38.4 | 38.4 | 38.4 | 38.5 |
| Germany | 0.0 | 0.0 | 0.0 | 0.0 | 35.4 | 35.3 | 35.1 | 34.9 | 34.6 | 34.2 | 33.8 | 33.3 | 32.8 | 32.3 |
| Greece | 0.0 | 0.0 | 0.0 | 0.0 | 0.0 | 4.8 | 4.7 | 4.6 | 4.6 | 4.5 | 4.5 | 4.5 | 4.4 | 4.4 |
| Hungary | 0.0 | 0.5 | 2.3 | 4.5 | 4.5 | 4.5 | 4.5 | 4.5 | 4.5 | 4.4 | 4.4 | 4.3 | 4.2 | 4.2 |
| Iceland | 0.0 | 0.1 | 0.1 | 0.2 | 0.2 | 0.2 | 0.2 | 0.2 | 0.2 | 0.2 | 0.2 | 0.2 | 0.2 | 0.2 |
| Ireland | 0.0 | 1.5 | 1.5 | 1.4 | 2.9 | 2.8 | 2.8 | 2.7 | 2.7 | 2.7 | 2.7 | 2.7 | 2.7 | 2.7 |
| Israel | 0.0 | 0.0 | 0.0 | 0.0 | 8.9 | 8.9 | 9.0 | 9.1 | 9.2 | 9.3 | 9.4 | 9.5 | 9.6 | 9.7 |
| Italy | 0.0 | 0.0 | 0.0 | 0.0 | 26.4 | 26.3 | 26.2 | 26.1 | 26.0 | 25.9 | 25.9 | 25.9 | 25.8 | 25.8 |
| Japan | 0.0 | 4.6 | 22.4 | 43.7 | 42.6 | 41.7 | 40.9 | 40.2 | 39.7 | 39.2 | 38.8 | 38.4 | 38.1 | 37.8 |
| Korea | 0.0 | 2.3 | 11.3 | 22.6 | 22.5 | 22.5 | 22.4 | 22.4 | 22.3 | 22.2 | 22.1 | 21.9 | 21.7 | 21.4 |
| Latvia | 0.0 | 0.0 | 0.0 | 0.0 | 0.0 | 1.1 | 1.1 | 1.1 | 1.0 | 1.0 | 1.0 | 1.0 | 0.9 | 0.9 |
| Luxembourg | 0.0 | 0.0 | 0.2 | 0.3 | 0.3 | 0.3 | 0.3 | 0.3 | 0.3 | 0.3 | 0.3 | 0.3 | 0.3 | 0.3 |
| Mexico | 0.0 | 11.0 | 55.1 | 110.0 | 109.9 | 109.8 | 109.6 | 109.4 | 109.2 | 109.0 | 108.7 | 108.4 | 108.1 | 107.8 |
| Netherlands | 0.0 | 0.9 | 4.7 | 9.3 | 9.4 | 9.4 | 9.4 | 9.4 | 9.5 | 9.5 | 9.5 | 9.4 | 9.4 | 9.4 |
| New Zealand | 0.0 | 0.0 | 0.0 | 0.0 | 3.1 | 3.1 | 3.1 | 3.1 | 3.1 | 3.2 | 3.2 | 3.2 | 3.2 | 3.2 |
| Norway | 0.0 | 0.3 | 1.5 | 3.0 | 3.0 | 3.0 | 3.1 | 3.1 | 3.1 | 3.1 | 3.1 | 3.1 | 3.1 | 3.1 |
| Poland | 0.0 | 0.0 | 0.0 | 0.0 | 0.0 | 18.1 | 17.7 | 17.2 | 16.7 | 16.2 | 15.8 | 15.4 | 15.0 | 14.6 |
| Portugal | 0.0 | 0.0 | 0.0 | 0.0 | 0.0 | 3.7 | 3.7 | 3.7 | 3.6 | 3.6 | 3.6 | 3.6 | 3.6 | 3.6 |
| Slovak Republic | 0.0 | 0.0 | 0.0 | 0.0 | 0.0 | 2.4 | 2.4 | 2.3 | 2.3 | 2.2 | 2.2 | 2.1 | 2.1 | 2.1 |
| Slovenia | 0.0 | 0.0 | 0.0 | 0.0 | 0.0 | 1.0 | 1.0 | 1.0 | 0.9 | 0.9 | 0.9 | 0.9 | 0.9 | 0.9 |
| Spain | 0.0 | 2.0 | 9.4 | 18.3 | 17.8 | 17.3 | 16.9 | 16.5 | 16.2 | 15.9 | 15.7 | 15.5 | 15.3 | 15.2 |
| Sweden | 0.0 | 0.6 | 3.1 | 6.3 | 6.4 | 6.4 | 6.5 | 6.5 | 6.4 | 6.4 | 6.3 | 6.2 | 6.2 | 6.1 |
| Switzerland | 0.0 | 0.4 | 1.9 | 3.7 | 3.8 | 3.8 | 3.7 | 3.7 | 3.7 | 3.7 | 3.7 | 3.7 | 3.6 | 3.6 |
| Turkey | 0.0 | 6.0 | 29.7 | 59.2 | 58.9 | 58.7 | 58.5 | 58.2 | 58.0 | 57.7 | 57.4 | 57.1 | 56.8 | 56.4 |
| United Kingdom | 4.0 | 20.0 | 20.0 | 40.2 | 40.2 | 40.2 | 40.1 | 40.0 | 39.9 | 39.8 | 39.7 | 39.6 | 39.5 | 39.4 |
| United States | 21.5 | 108.0 | 108.6 | 218.1 | 218.9 | 219.7 | 220.3 | 220.7 | 221.1 | 221.5 | 221.8 | 222.0 | 222.2 | 222.5 |
| Brazil | 0.0 | 13.0 | 64.1 | 127.2 | 126.4 | 125.6 | 124.9 | 124.2 | 123.5 | 122.6 | 121.6 | 120.4 | 119.0 | 117.5 |
| Bulgaria | 0.0 | 0.0 | 0.0 | 0.0 | 0.0 | 3.3 | 3.2 | 3.1 | 3.0 | 3.0 | 2.9 | 2.9 | 2.8 | 2.8 |
| China (People's Republic of) | 0.0 | 443.4 | 434.8 | 425.5 | 824.0 | 802.9 | 785.8 | 770.5 | 754.9 | 740.5 | 729.0 | 719.6 | 711.8 | 705.5 |
| Cyprus | 0.0 | 0.0 | 0.0 | 0.0 | 0.0 | 0.6 | 0.6 | 0.6 | 0.6 | 0.6 | 0.6 | 0.6 | 0.6 | 0.6 |
| India | 0.0 | 121.8 | 603.7 | 1200.1 | 1191.2 | 1187.1 | 1186.5 | 1185.8 | 1182.8 | 1181.7 | 1177.0 | 1169.7 | 1161.6 | 1154.4 |
| Indonesia | 0.0 | 0.0 | 0.0 | 0.0 | 0.0 | 223.2 | 221. | 219.4 | 218.4 | 217.8 | 217.6 | 217.8 | 218.0 | 217.8 |
|  |  |  |  |  |  |  | 4 |  |  |  |  |  |  |  |
| Lithuania | 0.0 | 0.0 | 0.0 | 0.0 | 0.0 | 1.5 | 1.5 | 1.5 | 1.5 | 1.5 | 1.5 | 1.4 | 1.4 | 1.4 |
| Malta | 0.0 | 0.0 | 0.1 | 0.2 | 0.2 | 0.2 | 0.2 | 0.2 | 0.2 | 0.2 | 0.2 | 0.2 | 0.2 | 0.2 |
| Romania | 0.0 | 0.0 | 0.0 | 0.0 | 0.0 | 10.1 | 10.0 | 9.8 | 9.7 | 9.5 | 9.4 | 9.3 | 9.2 | 9.1 |
| Russia | 0.0 | 9.4 | 46.6 | 91.4 | 89.4 | 87.2 | 85.1 | 83.0 | 80.9 | 78.9 | 77.0 | 75.2 | 73.7 | 72.5 |
| South Africa | 0.0 | 5.1 | 25.6 | 51.1 | 51.2 | 50.9 | 50.2 | 49.6 | 49.1 | 48.6 | 48.3 | 48.2 | 48.1 | 48.0 |

Half the population of G20 countries will, at some point in their lives, be diagnosed with a **cancer** event. Depending on the utility of genomics for informing the treatment of different cancer types, healthcare systems could be deploying cancer genomics for over 40 million people in the developed world by 2025 (see Table 3).

**Table 3.**
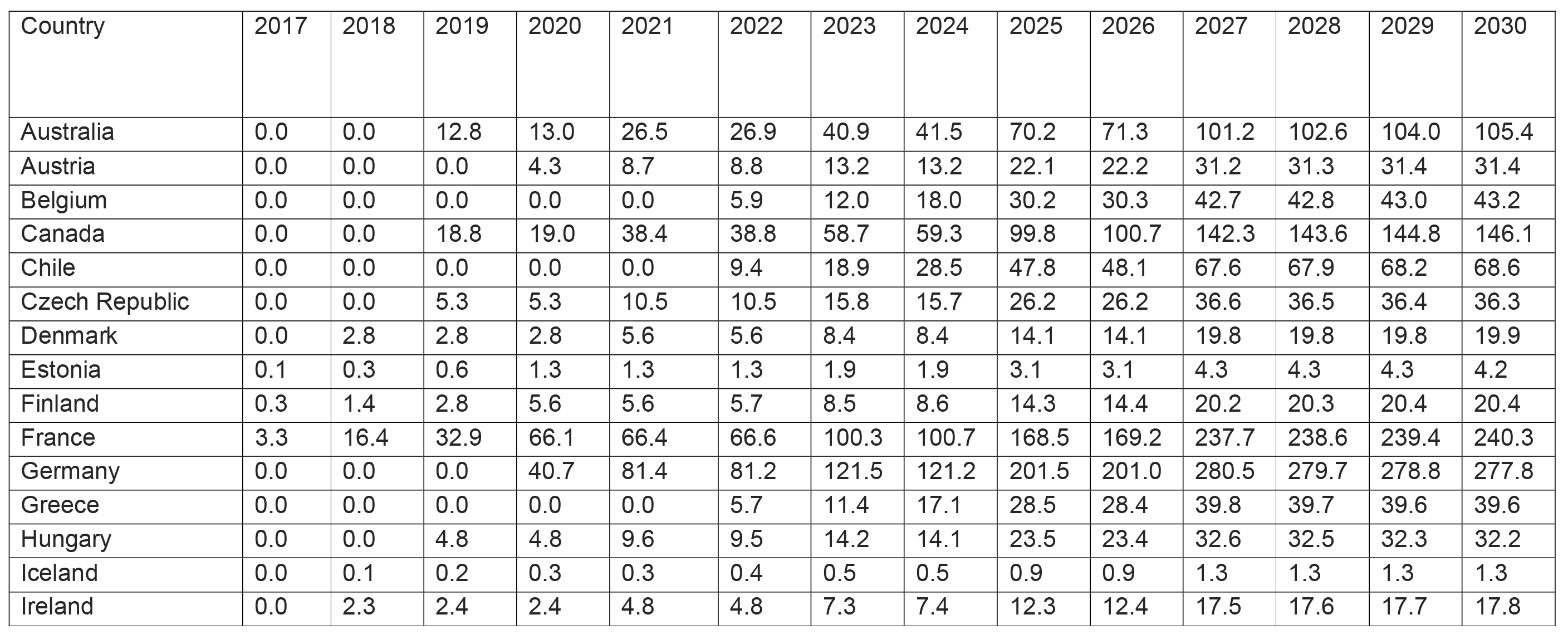
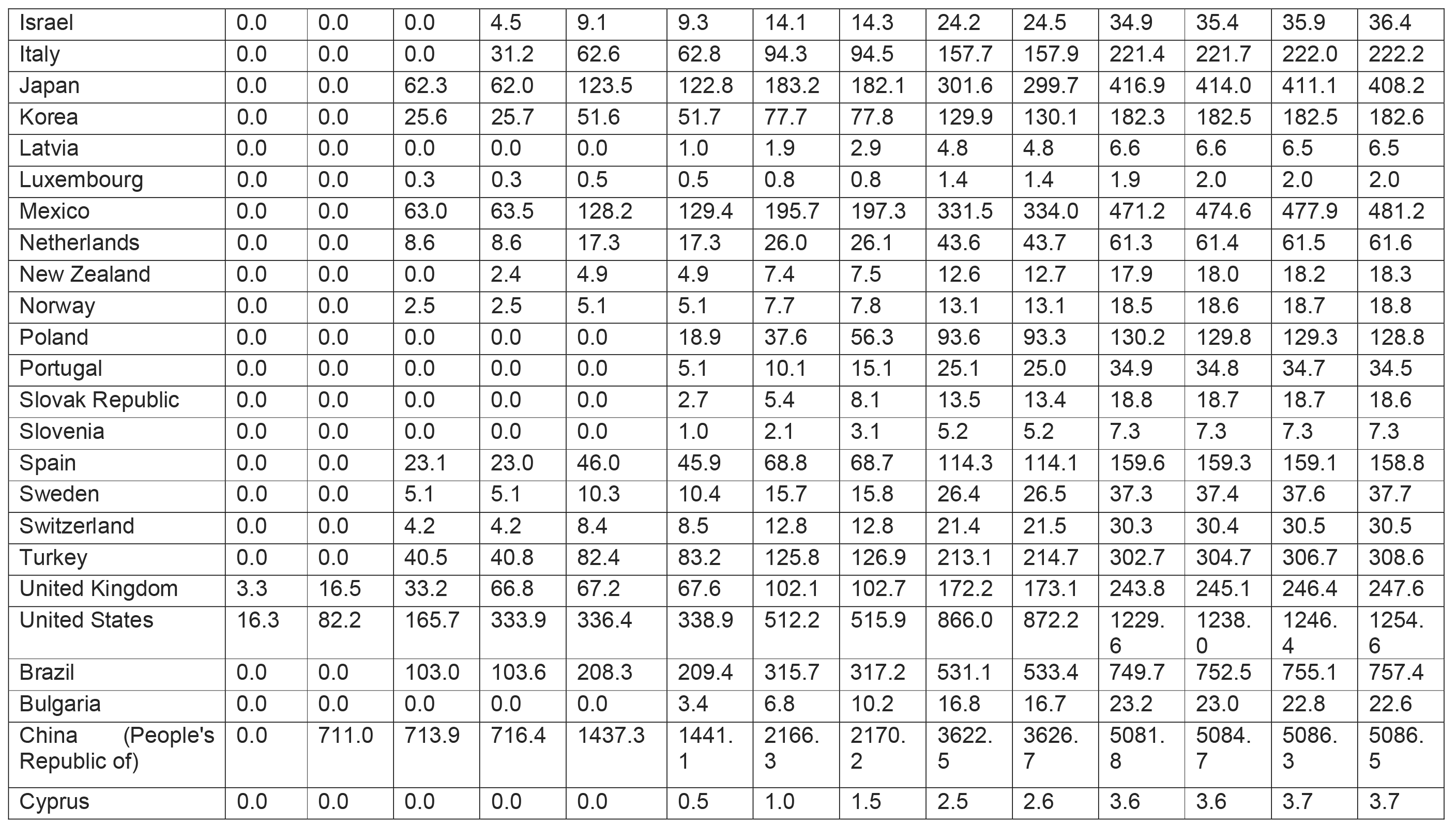
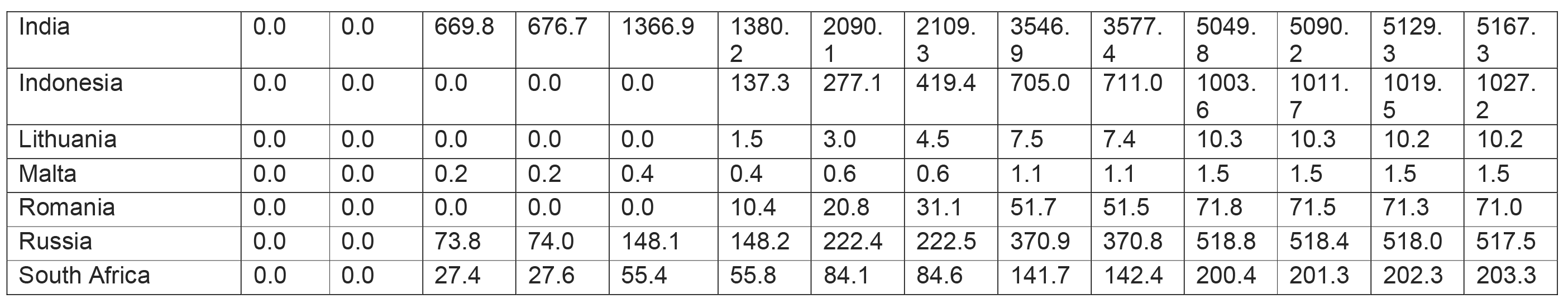
Estimated number of Cancer genomes (in thousands) sequenced in each OECD country over time

| Country | 2017 | 2018 | 2019 | 2020 | 2021 | 2022 | 2023 | 2024 | 2025 | 2026 | 2027 | 2028 | 2029 | 2030 |
| --- | --- | --- | --- | --- | --- | --- | --- | --- | --- | --- | --- | --- | --- | --- |
| Australia | 0.0 | 0.0 | 12.8 | 13.0 | 26.5 | 26.9 | 40.9 | 41.5 | 70.2 | 71.3 | 101.2 | 102.6 | 104.0 | 105.4 |
| Austria | 0.0 | 0.0 | 0.0 | 4.3 | 8.7 | 8.8 | 13.2 | 13.2 | 22.1 | 22.2 | 31.2 | 31.3 | 31.4 | 31.4 |
| Belgium | 0.0 | 0.0 | 0.0 | 0.0 | 0.0 | 5.9 | 12.0 | 18.0 | 30.2 | 30.3 | 42.7 | 42.8 | 43.0 | 43.2 |
| Canada | 0.0 | 0.0 | 18.8 | 19.0 | 38.4 | 38.8 | 58.7 | 59.3 | 99.8 | 100.7 | 142.3 | 143.6 | 144.8 | 146.1 |
| Chile | 0.0 | 0.0 | 0.0 | 0.0 | 0.0 | 9.4 | 18.9 | 28.5 | 47.8 | 48.1 | 67.6 | 67.9 | 68.2 | 68.6 |
| Czech Republic | 0.0 | 0.0 | 5.3 | 5.3 | 10.5 | 10.5 | 15.8 | 15.7 | 26.2 | 26.2 | 36.6 | 36.5 | 36.4 | 36.3 |
| Denmark | 0.0 | 2.8 | 2.8 | 2.8 | 5.6 | 5.6 | 8.4 | 8.4 | 14.1 | 14.1 | 19.8 | 19.8 | 19.8 | 19.9 |
| Estonia | 0.1 | 0.3 | 0.6 | 1.3 | 1.3 | 1.3 | 1.9 | 1.9 | 3.1 | 3.1 | 4.3 | 4.3 | 4.3 | 4.2 |
| Finland | 0.3 | 1.4 | 2.8 | 5.6 | 5.6 | 5.7 | 8.5 | 8.6 | 14.3 | 14.4 | 20.2 | 20.3 | 20.4 | 20.4 |
| France | 3.3 | 16.4 | 32.9 | 66.1 | 66.4 | 66.6 | 100.3 | 100.7 | 168.5 | 169.2 | 237.7 | 238.6 | 239.4 | 240.3 |
| Germany | 0.0 | 0.0 | 0.0 | 40.7 | 81.4 | 81.2 | 121.5 | 121.2 | 201.5 | 201.0 | 280.5 | 279.7 | 278.8 | 277.8 |
| Greece | 0.0 | 0.0 | 0.0 | 0.0 | 0.0 | 5.7 | 11.4 | 17.1 | 28.5 | 28.4 | 39.8 | 39.7 | 39.6 | 39.6 |
| Hungary | 0.0 | 0.0 | 4.8 | 4.8 | 9.6 | 9.5 | 14.2 | 14.1 | 23.5 | 23.4 | 32.6 | 32.5 | 32.3 | 32.2 |
| Iceland | 0.0 | 0.1 | 0.2 | 0.3 | 0.3 | 0.4 | 0.5 | 0.5 | 0.9 | 0.9 | 1.3 | 1.3 | 1.3 | 1.3 |
| Ireland | 0.0 | 2.3 | 2.4 | 2.4 | 4.8 | 4.8 | 7.3 | 7.4 | 12.3 | 12.4 | 17.5 | 17.6 | 17.7 | 17.8 |
| Israel | 0.0 | 0.0 | 0.0 | 4.5 | 9.1 | 9.3 | 14.1 | 14.3 | 24.2 | 24.5 | 34.9 | 35.4 | 35.9 | 36.4 |
| Italy | 0.0 | 0.0 | 0.0 | 31.2 | 62.6 | 62.8 | 94.3 | 94.5 | 157.7 | 157.9 | 221.4 | 221.7 | 222.0 | 222.2 |
| Japan | 0.0 | 0.0 | 62.3 | 62.0 | 123.5 | 122.8 | 183.2 | 182.1 | 301.6 | 299.7 | 416.9 | 414.0 | 411.1 | 408.2 |
| Korea | 0.0 | 0.0 | 25.6 | 25.7 | 51.6 | 51.7 | 77.7 | 77.8 | 129.9 | 130.1 | 182.3 | 182.5 | 182.5 | 182.6 |
| Latvia | 0.0 | 0.0 | 0.0 | 0.0 | 0.0 | 1.0 | 1.9 | 2.9 | 4.8 | 4.8 | 6.6 | 6.6 | 6.5 | 6.5 |
| Luxembourg | 0.0 | 0.0 | 0.3 | 0.3 | 0.5 | 0.5 | 0.8 | 0.8 | 1.4 | 1.4 | 1.9 | 2.0 | 2.0 | 2.0 |
| Mexico | 0.0 | 0.0 | 63.0 | 63.5 | 128.2 | 129.4 | 195.7 | 197.3 | 331.5 | 334.0 | 471.2 | 474.6 | 477.9 | 481.2 |
| Netherlands | 0.0 | 0.0 | 8.6 | 8.6 | 17.3 | 17.3 | 26.0 | 26.1 | 43.6 | 43.7 | 61.3 | 61.4 | 61.5 | 61.6 |
| New Zealand | 0.0 | 0.0 | 0.0 | 2.4 | 4.9 | 4.9 | 7.4 | 7.5 | 12.6 | 12.7 | 17.9 | 18.0 | 18.2 | 18.3 |
| Norway | 0.0 | 0.0 | 2.5 | 2.5 | 5.1 | 5.1 | 7.7 | 7.8 | 13.1 | 13.1 | 18.5 | 18.6 | 18.7 | 18.8 |
| Poland | 0.0 | 0.0 | 0.0 | 0.0 | 0.0 | 18.9 | 37.6 | 56.3 | 93.6 | 93.3 | 130.2 | 129.8 | 129.3 | 128.8 |
| Portugal | 0.0 | 0.0 | 0.0 | 0.0 | 0.0 | 5.1 | 10.1 | 15.1 | 25.1 | 25.0 | 34.9 | 34.8 | 34.7 | 34.5 |
| Slovak Republic | 0.0 | 0.0 | 0.0 | 0.0 | 0.0 | 2.7 | 5.4 | 8.1 | 13.5 | 13.4 | 18.8 | 18.7 | 18.7 | 18.6 |
| Slovenia | 0.0 | 0.0 | 0.0 | 0.0 | 0.0 | 1.0 | 2.1 | 3.1 | 5.2 | 5.2 | 7.3 | 7.3 | 7.3 | 7.3 |
| Spain | 0.0 | 0.0 | 23.1 | 23.0 | 46.0 | 45.9 | 68.8 | 68.7 | 114.3 | 114.1 | 159.6 | 159.3 | 159.1 | 158.8 |
| Sweden | 0.0 | 0.0 | 5.1 | 5.1 | 10.3 | 10.4 | 15.7 | 15.8 | 26.4 | 26.5 | 37.3 | 37.4 | 37.6 | 37.7 |
| Switzerland | 0.0 | 0.0 | 4.2 | 4.2 | 8.4 | 8.5 | 12.8 | 12.8 | 21.4 | 21.5 | 30.3 | 30.4 | 30.5 | 30.5 |
| Turkey | 0.0 | 0.0 | 40.5 | 40.8 | 82.4 | 83.2 | 125.8 | 126.9 | 213.1 | 214.7 | 302.7 | 304.7 | 306.7 | 308.6 |
| United Kingdom | 3.3 | 16.5 | 33.2 | 66.8 | 67.2 | 67.6 | 102.1 | 102.7 | 172.2 | 173.1 | 243.8 | 245.1 | 246.4 | 247.6 |
| United States | 16.3 | 82.2 | 165.7 | 333.9 | 336.4 | 338.9 | 512.2 | 515.9 | 866.0 | 872.2 | 1229.6 | 1238.0 | 1246.4 | 1254.6 |
| Brazil | 0.0 | 0.0 | 103.0 | 103.6 | 208.3 | 209.4 | 315.7 | 317.2 | 531.1 | 533.4 | 749.7 | 752.5 | 755.1 | 757.4 |
| Bulgaria | 0.0 | 0.0 | 0.0 | 0.0 | 0.0 | 3.4 | 6.8 | 10.2 | 16.8 | 16.7 | 23.2 | 23.0 | 22.8 | 22.6 |
| China (People's Republic of) | 0.0 | 711.0 | 713.9 | 716.4 | 1437.3 | 1441.1 | 2166.3 | 2170.2 | 3622.5 | 3626.7 | 5081.8 | 5084.7 | 5086.3 | 5086.5 |
| Cyprus | 0.0 | 0.0 | 0.0 | 0.0 | 0.0 | 0.5 | 1.0 | 1.5 | 2.5 | 2.6 | 3.6 | 3.6 | 3.7 | 3.7 |
| GA4GH Connect, October 2017 |  |  |  |  |  |  |  |  |  |  |  |  |  |  |
| India | 0.0 | 0.0 | 669.8 | 676.7 | 1366.9 | 1380.2 | 2090.1 | 2109.3 | 3546.9 | 3577.4 | 5049.8 | 5090.2 | 5129.3 | 5167.3 |
| Indonesia | 0.0 | 0.0 | 0.0 | 0.0 | 0.0 | 137.3 | 277.1 | 419.4 | 705.0 | 711.0 | 1003.6 | 1011.7 | 1019.5 | 1027.2 |
| Lithuania | 0.0 | 0.0 | 0.0 | 0.0 | 0.0 | 1.5 | 3.0 | 4.5 | 7.5 | 7.4 | 10.3 | 10.3 | 10.2 | 10.2 |
| Malta | 0.0 | 0.0 | 0.2 | 0.2 | 0.4 | 0.4 | 0.6 | 0.6 | 1.1 | 1.1 | 1.5 | 1.5 | 1.5 | 1.5 |
| Romania | 0.0 | 0.0 | 0.0 | 0.0 | 0.0 | 10.4 | 20.8 | 31.1 | 51.7 | 51.5 | 71.8 | 71.5 | 71.3 | 71.0 |
| Russia | 0.0 | 0.0 | 73.8 | 74.0 | 148.1 | 148.2 | 222.4 | 222.5 | 370.9 | 370.8 | 518.8 | 518.4 | 518.0 | 517.5 |
| South Africa | 0.0 | 0.0 | 27.4 | 27.6 | 55.4 | 55.8 | 84.1 | 84.6 | 141.7 | 142.4 | 200.4 | 201.3 | 202.3 | 203.3 |

**Common diseases** affect nearly everyone. When genomic information can be used clinically for broad, common diseases, it will be feasible to justify sequencing entire populations. Whole-population-scale sequencing is feasible currently for smaller countries (and is largely in place already in some countries, e.g. Iceland), and is likely to be in place at some point in the next two decades.

We developed a crude model for estimating the rate of healthcare-funded genomes worldwide. This accounts for the preparedness of each country for genomics, and infers a progression of cancer-related sequencing over time (conservative estimate: genomics is used in 70% of cancer diagnoses by 2027). Figure 1 shows the growth in the number of genomes based on this model, with an expected number of 47.5 million genomes sequenced for rare disease diagnoses (patients + family members) and 83 million (cancer + matched normal) genome sequences for cancer by 2025.

**Figure 1.**
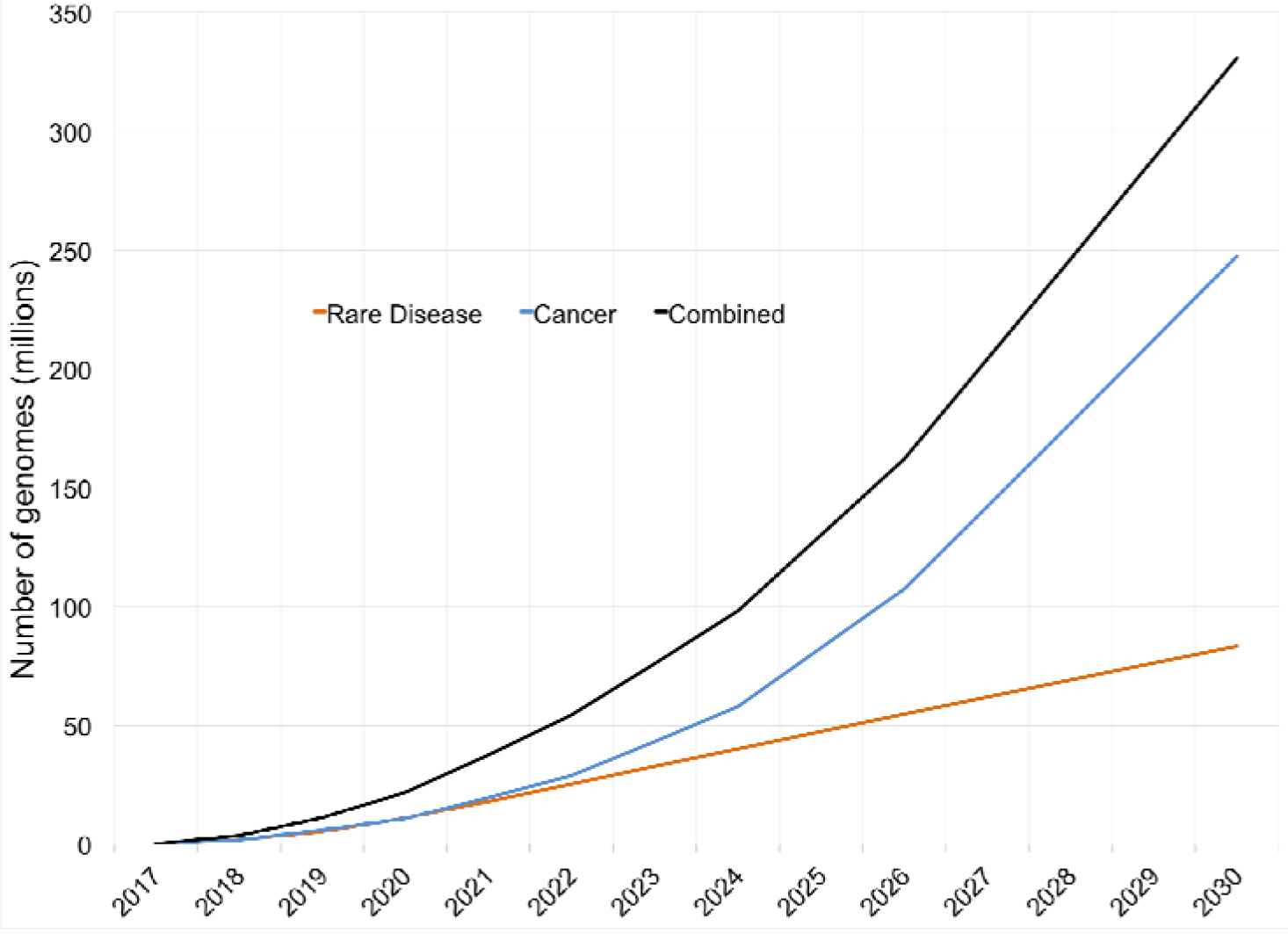
The number of genomes sequenced per rare disease proband was assumed to be 2.3, with 1% of young children estimated to be affected by a rare disease. Due to the complexities of applying cancer genomics in the clinic, we estimate a slower rate of increase in healthcare-funded cancer genomes with a ‘ramp up’ in numbers starting from 2022.

This model is crude in its treatment of the complex, policy heavy area of genomic medicine, and undoubtedly some countries will have a faster or slower uptake than we predict. Furthermore, if the cost of sequencing falls radically, full population-scale sequencing may be deployed quite quickly, and so complete population sequencing could occur in this time frame.

Whatever the details of the country-by-country roll out, we are confident that over 60 million patients will have their genome sequenced in a healthcare context by 2025. This represents a large technical challenge for healthcare systems, and a huge opportunity for research.

## The need for cohorts of 10 million+ humans

The availability of large cohorts that represent individuals from different populations throughout the world would be transformative for human research. The advantages of large cohorts are considerable, especially when the costs of raw sequencing and first-line analysis will be borne primarily by healthcare systems all over the world. This could be seen as having spill-over benefits for research, which generally functions within a narrow economy. The use of healthcare-driven genomic data being also repositioned for research is part of the broad trend of breaking down barriers between research and healthcare, providing for a more permeable, ‘learning’ healthcare system with deep, sustained ties to research.

The research benefits that accrue from assembling cohorts of over 1 million people (currently the largest cohorts available) are an open question. Cohorts of 0.5 to 1 million people might be adequate for research, and tracking more than that may not be necessary. However, experience has shown us that larger sample sizes allow for more fine-grained analysis (12) and more robust findings in genetics, regardless of the system used to collect them.

We have identified eight major practical, principled arguments to support our position that virtual cohorts of 100 million people or more would have tangible research benefits.

### 1. Robust statistical association of rare and extremely rare events

Many rare diseases occur at person frequencies of 1 every 10,000 to 1 every 1,000,000 live births. To associate a particular mutation to a particular phenotype robustly, at least two observations are required, often more. This argues for pooling rare disease diagnoses in populations of 10 million to 500 million, where all the suspected rare disease individuals (between 1% to 2%) and their parents (where available) are sequenced.

The Matchmaker Exchange (MME) project (13) is building a platform for matching patients with similar phenotypic and genotypic profiles through standardized application programming interfaces (APIs) and procedural conventions. Its participating genomic groups are located in multiple countries, and pool possible diagnoses with the goal of finding ‘a second match’. MME has produced nearly 30 matches as of October 2017, predominantly matching patients who live in different countries (i.e. not enrolled in the same healthcare system).

### 2. Robust assessment of penetrance of disease or trait-causing alleles

The penetrance of potential disease-causing alleles is best assessed using broadly ascertained (e.g. population-scale) cohorts. Genomics of rare disease (patients and their parents) and the ‘healthy’ genomes of cancer patients can be deployed effectively to understand the frequency of putative alleles, irrespective of the specific disease of interest.

### 3. Ability to query very low frequency alleles in specific environments

Many phenotypically interesting alleles, in particular disabling mutations such as stop codons, are at low frequency (i.e. below 0.1%). Many diseases only occur in, or are at higher frequency in, specific environments. To observe relevant shifts in the ratios of disease, sample sizes of between 1 million and 10 million are necessary.

Thanks to the ease of interpretation of some of these rare alleles, in particular the clear-cut loss-of-function alleles, both positive association (i.e. clear association of an allele with a disease) and negative association are of great interest to the pharmaceutical and biotech industries.

### 4. Heterogeneous diseases such as cancer benefit from large sample size

Considering the combinatorics of all possible mutations in all tissue types in an array of environmental conditions (e.g. immune status of the patient), each cancer is best thought of as a unique, uncontrolled somatic aberration. Large sample sizes make taming this heterogeneity more feasible, as they allow robust statistical analysis and rigorous testing of the resulting models in data not used to train models.

### 5. Discovering and characterising epistasis is more feasible with large sample sizes

Epistasis (i.e. non-linear interactions between alleles) is commonplace in model-organism genetics. It is difficult to utilise in human genetics, principally because interesting alleles are low frequency and the interaction occurs as the product of these frequencies; however, one-million-person or more cohorts overcomes these frequency concerns and provides a starting point for probing these interactions. Each order of magnitude higher will improve discoverability of rarer epistatic events. Such discoveries need not account for whole-population-level variance to be interesting; they are inherently useful for understanding the molecular biology of a disease or trait of interest.

### 6. Large, diverse cohorts provide a baseline understanding of the selection pressures on every base in the human genome

When we reach the ‘10 million to 100 million genomes sequenced’ milestone, we will be in the final approach to the 1.1e-8 mutation rate for each base in the human genome. Even at the current cohort size of around 100,000 in the ExAC and gnomAD databases, we can derive a reasonably robust model for mutation rates and gene-scale selection. With higher sample numbers, these models can become more sophisticated in terms of the heterogeneity in mutation rates they model, and narrower in terms of the genomic region they probe—ultimately to single-base-pair resolution.

### 7. Very large cohorts are ideal for Mendelian randomisation approaches in understanding causality in observational studies

Germline genetic information has two unique properties that make it invaluable for observational (epidemiological) studies: stability of germline variation and randomisation of allele distribution in a population. Germline variation rarely changes over the lifetime of an individual. If it does, it is readily detected in the form of mosaicism or cancer. Variants are randomly assigned to chromosomes during meiosis, and this randomisation is present in the resulting distribution of alleles inside a population.

Genetic variation can be used as a subtle ‘natural experiment’ to clarify the relationships between variable measures (e.g. weight gain and Type II diabetes, or alcohol usage during pregnancy and outcome of offspring). The genetic variables act as unbiased statistical instruments to provide insight on causality. These techniques work with very-low-impact variants, but require large sample sizes (millions of individuals) to work well.

### 8. Variation in allele frequency, environments, and measurements worldwide can be leveraged

The precise allele frequencies present in the germline vary between populations, mainly due to population drift accentuated by bottlenecks. Different physical and social environments influence environmental exposures and developmental trajectories, and different healthcare systems vary in their ability to measure specific diseases and traits.

At scales below 10,000 individuals, these differences hinder joint use of datasets and force researchers to focus only on the larger subsets present in their data. At larger scales, this diversity can be leveraged to find more conclusive evidence behind specific alleles and environmental differences. It also opens the door to serendipitous discoveries based on traits that are measured well by one system and not another.

## Challenges

### Ethical and regulatory challenges

Ethical consideration of patients and populations together with responsible regulation are critical to biomedical research in all its forms. This is particularly true for healthcare-funded genomics, which involves deeply complex national regulation and legislation. Keeping the UN *Universal Declaration of Human Rights* at the heart of these conversations will serve to activate the right ‘to share in scientific advancement and its benefits’ (14), firmly rooted in concerns shared by people of all nations. It also ensures a common human-rights approach to addressing the benefits and potential risks in a balanced manner.

Each society has its own, unique perspective on the sharing of personal information, with more open or restrictive regulatory norms and systems on data collection, access, and sharing. The commonality is that all of these systems have some mechanism by which researchers are able to access both research and clinical data, specifically to use it in ways that enable patients and research participants to share in scientific advancement and its benefits.

Population-scale sequencing schemes—wherein regulated healthcare providers share clinical genomic data for research—are unlikely to allow large-scale aggregation of data to migrate beyond national boundaries, but it is feasible to imagine that federated analysis without data movement is possible. Accountability principles (15) and sanctions for misuse are being developed in order to respect and maintain the trust of participants.

Based on the universal principles of benefitting from scientific research and respecting national regulatory architecture, we believe that most healthcare systems can ultimately participate in responsible, worldwide data sharing while remaining compliant with applicable jurisdictional law and institutional policies.

### Technical challenges

To realise the goal of deploying single, agreed methods in multiple locations offering harmonised datasets, each location must present genomic and associated phenotypic information in a consistent, standard manner. These standards should, wherever possible, be based on a service architecture in which information is retrieved using web-based calls (e.g. REST protocol). This is the de facto standard for large-scale data delivery in science and technology.

Currently, genomics data analysis is tied to bespoke, file-based systems, with a mixture of domain-specific standard file types tied together in institutional (and sometimes individual researcher) configurations. This scheme is incompatible with a global, federated architecture. A major goal of GA4GH is to provide diverse, service-based standards to enable a global federation of identity and data, within a robust, security-assured cloud environment that enforces jurisdictional constraints and local service agreements.

Researcher access to genomic data refers specifically to access via analytic machinery created for research users. Over the past decade there has been a significant shift towards virtualisation in computing. This involves packaging and distributing the computing components (e.g. processors, storage units, platforms) or entire computing infrastructures and allowing them to run in an environment that simulates a specific computational architecture as if it were physically co-located.

The value of virtualisation is that it enables a third-party service provider (“cloud” provider) to provision computing services and storage capacity “on demand” as they are needed. “Cloud” services are particularly well suited to computing problems involving large volumes of storage and complex computations (e.g. genomic research). Virtualisation is now deployed on a large scale in cloud environments, enabling researchers to use virtual software tools to analyse data distributed across multiple physical data stores, rather than downloading large amounts of data for use with local software tools. This pattern of moving the analysis to the data rather than data to the analysis allows for more appropriate scaling and control of access via national schemes, while also enabling consistency of analysis by a researcher “visiting” each of these schemes.

Another technology trend that holds great potential for genomics research is federation— both identity federation and data federation. Identity federation enables multiple organisations to rely on each other or a third party to authenticate researchers’ identities and to share security attributes. This enables each data holder to enforce its own security policies, using attributes passed along with the authenticated identity. Data federation enables genomic analysis to be performed across multiple data stores.

### Security challenges

GA4GH, as an international consortium federating large volumes of sensitive clinical and genomic data across virtual computing environments, is presented with formidable challenges in assuring data confidentiality, data integrity, service availability, and individual privacy. Some of these challenges call for innovative application of well-established security standards, frameworks, and protocols, such as identity federation on a global scale. Other challenges require solutions still emerging from security research, such as privacy-preserving data linkage and homomorphic encryption. GA4GH seeks to apply current and emerging technology solutions, standards, and best practices to help protect the confidentiality, integrity, and availability of sensitive, high-value data. Healthcare data are a leading target for cyber-attackers; as such, GA4GH and its partners must implement a layered and proactive scheme to identify potential threats and vulnerabilities, continuously monitor the use of data and services, detect potential attacks, and collectively respond to potential breaches. Risk management is central to GA4GH’s standards-development process, while seeking to leverage industry standards and best practices wherever possible.

### Societal challenges

Ethical, regulatory, and technical challenges all feed into the wider societal challenges of bringing genomics into healthcare, which we must meet as a single community. The three strands of this challenge are: maintaining public trust in healthcare systems, overcoming differences in objectives and methods between research and healthcare, and breaking down unproductive divides between disciplines.

Our future vision of healthcare is one in which vetted researchers around the world can, with appropriate oversight and policy enforcement, gain access to anonymised human health data. This system of access is based on trust: of researchers, institutions, countries, and participants—similar to the trust on which current global biomedical research depends. Local, national, and global dialogue among researchers, clinicians, and participants is active, engaged, and useful. Trust in this paradigm is earned and maintained by respectful, open discussion and collaboration among all parties.

We envision clinical and basic researchers collaborating seamlessly in the context of practising healthcare. The virtuous cycle of research and healthcare will be celebrated as an example of harmonised human endeavour. The track record of data science makes these ambitions realistic. Genomic datasets from healthcare will be among the largest generated over the next decade, and to make the most of them we need to harness the best of computational biology: both academic (i.e. electrical engineering, computer science, statistics, physics) and commercial (i.e. technology, pharmaceutical, biotech, health informatics).

Communication and respect for all players will maintain a steady focus on shared objectives and outcomes.

## What if there were no standards?

A sceptic might wonder whether this is really needed. Perhaps large healthcare systems have enough internal drivers to ensure good delivery of genomic medicine without any global coordination, and smaller healthcare systems will naturally align themselves to their nearest scheme (culturally or physically). Perhaps researchers are better off negotiating individually, system by system, hospital by hospital, thus allowing more local innovation. Do we really need to coordinate worldwide to deliver dividends on genomics?

We have already established that we need to coordinate between large healthcare systems in rare-disease diagnosis and discovery. As a practical example, Matchmaker Exchange illustrates the power of bringing practicing clinicians and researchers together. This is also mirrored in the Cancer field where large international projects have switched to using more routine healthcare generated data sources.

Furthermore, as nascent genomic medicine schemes are being delivered in a variety of countries, it is clear that the federated approach enabled by GA4GH is the only scheme that can satisfy both research and healthcare goals. In addition, many commercial and public organizations want to minimize the costs and risks of the complex technical software needed to either contribute to genomic medicine or deliver genomic tools. A complex, multistakeholder ecosystem requires neutral and technically competent standards.

Nevertheless, standards and frameworks must be fit for purpose and useful for the broad set of users: clinical, academic, commercial, and public. These standards must also enable progress—not stifle it. To these ends, GA4GH is focused on the creation and management of genomics standards and not on their implementations, though we expect and encourage many implementing groups to actively participate.

## Conclusion

It is not difficult to imagine a world in which genomic and molecular data have been collected for over a billion humans. In such a future, genomics will deliver both personalised and population-based benefits to patients—in acute healthcare, preventative medicine and probably unforeseen ways. When researchers everywhere can access secure data responsibly and use the data to benefit human health, discoveries will accelerate in both health and fundamental biology, in areas on which future benefits may unexpectedly rely.

These goals are as ambitious in scope as the coordinated, international response to infectious disease outbreaks, and the international network of seismic sensors for early detection of earthquakes and tsunamis—both of which are demonstrably successful. To achieve this ambition we must have ongoing, sustained public support, from individuals, organisations, and governments.

Just as the W3C consortium provides the framework for setting the standards of the World Wide Web, GA4GH has taken responsibility for building the technical standards—rooted in a robust regulatory, ethical, and security-assured framework—to enable healthcare to benefit directly from scientific progress. It represents an open culture that spans technical standards development and implementation, and discussions that inform and are informed by public policy.

Because the future of healthcare belongs to all of us, we warmly welcome the participation, on every level, of people in all disciplines, in every country. Built collaboratively by patients, clinicians, researchers, engineers, and their advocates, the future of healthcare could truly benefit all of humanity.

## Acknowledgements

We are grateful to members of GA4GH and the GA4GH Strategic Advisory Board for their feedback. Thanks to Angela Page and Mary Todd Bergman for help preparing this manuscript.

## Footnotes

1 For those deeply involved in genomic research, ‘genomics’ has a narrow meaning: assaying genomic DNA. However, the word has come to mean many things to the broader community. ‘Genomics’ encompasses molecular measurements (DNA, RNA, protein, and metabolites), subsequent data management and molecular data analysis. Each of these areas has its own specific discipline: transcriptomics, proteomics, metabolomics, bioinformatics. For those working alongside genomics, acknowledging these distinctions explicitly is clunky. New terms like ‘multi-omics’, rather than resolving the problem, create still more language barriers between research and clinical practise. In the interests of practicality and accuracy, we use the term ‘genomics’ to mean molecular, ‘genome-wide’ assays and their associated data management and analysis.

